# Unification of the M/ORF3-related proteins points to a diversified role for ion conductance in pathogenesis of coronaviruses and other nidoviruses

**DOI:** 10.1101/2020.11.10.377366

**Authors:** Yongjun Tan, Theresa Schneider, Prakash K. Shukla, Mahesh B. Chandrasekharan, L Aravind, Dapeng Zhang

## Abstract

The new coronavirus, SARS-CoV-2, responsible for the COVID-19 pandemic has emphasized the need for a better understanding of the evolution of virus-host conflicts. ORF3a in both SARS-CoV-1 and SARS-CoV-2 are ion channels (viroporins) and involved in virion assembly and membrane budding. Using sensitive profile-based homology detection methods, we unify the SARS-CoV ORF3a family with several families of viral proteins, including ORF5 from MERS-CoVs, proteins from beta-CoVs (ORF3c), alpha-CoVs (ORF3b), most importantly, the Matrix (M) proteins from CoVs, and more distant homologs from other nidoviruses. By sequence analysis and structural modeling, we show that these viral families utilize specific conserved polar residues to constitute an ion-conducting pore in the membrane. We reconstruct the evolutionary history of these families, objectively establish the common origin of the M proteins of CoVs and Toroviruses. We show that the divergent ORF3a/ORF3b/ORF5 families represent a duplication stemming from the M protein in alpha- and beta-CoVs. By phyletic profiling of major structural components of primary nidoviruses, we present a model for their role in virion assembly of CoVs, ToroVs and Arteriviruses. The unification of diverse M/ORF3 ion channel families in a wide range of nidoviruses, especially the typical M protein in CoVs, reveal a conserved, previously under-appreciated role of ion channels in virion assembly, membrane fusion and budding. We show that the M and ORF3 are under differential evolutionary pressures; in contrast to the slow evolution of M as core structural component, the CoV-ORF3 clade is under selection for diversification, which indicates it is likely at the interface with host molecules and/or immune attack.

**IMPORTANCE:** Coronaviruses (CoVs) have become a major threat to human welfare as the causative agents of several severe infectious diseases, namely Severe Acute Respiratory Syndrome (SARS), Middle Eastern Respiratory Syndrome (MERS), and the recently emerging human coronavirus disease 2019 (COVID-19). The rapid spread, severity of these diseases, as well as the potential re-emergence of other CoV-associated diseases have imposed a strong need for a thorough understanding of function and evolution of these CoVs. By utilizing robust domain-centric computational strategies, we have established homologous relationships between many divergent families of CoV proteins, including SARS-CoV/SARS-CoV-2 ORF3a, MERS-CoV ORF5, proteins from both beta-CoVs (ORF3c) and alpha-CoVs (ORF3b), the typical CoV Matrix proteins, and many distant homologs from other nidoviruses. We present evidence that they are active ion channel proteins, and the Cov-specific ORF3 clade proteins are under selection for rapid diversification, suggesting they might have been involved in interfering host molecules and/or immune attack.

## INTRODUCTION

The recent outbreak of human coronavirus disease 2019 (COVID-19) has generated a global health crisis (1). It is the 7^th^ human disease caused by coronaviruses, after Severe Acute Respiratory Syndrome (SARS) in 2003 (2), Middle Eastern Respiratory Syndrome (MERS) in 2012 (3), and four less-severe infections caused by human coronaviruses 229E (hCoV-229E) (4), hCoV-NL63 in 2004 (5), and hCoV-HKU1 in 2004 (6). Of these, SARS-CoV-2, SARS-CoV/SARS-CoV-1, MERS-CoV, hCoV-OC43 and hCoV-HKU1 belong to the beta coronavirus clade while hCoV-229E and hCoV-NL63 belong to the alpha coronavirus clade. Although the broad genomic structure and core gene-composition of these viruses is similar, the pathology and severity of these viruses, including SARS-CoV-2, are markedly distinct. According to the WHO report, as of October 2020, there have been over 360 million of confirmed cases with over 1 million deaths from COVID-19 globally. Therefore, the need for a better understanding of the biology and evolution of SARS-CoV-2 is a major desideratum to combat and prevent the disease.

Coronaviruses possess a large positive-sense single-stranded RNA genome with two third of the genome coding for the ORF1a/ORF1ab polyprotein. This is followed by several ORFs encoding so-called structural and accessory proteins, several of which might be variable between viruses (2). The ORF1 polyprotein is processed into smaller proteins by proteolytic cleavage catalyzed by one of its constituent components (the peptidase domain) (7), viral replication, viral RNA-processing (e.g. xEndoU endoRNase domain) and countering of defenses centered on NAD+/ADP-ribose (Macro domains) (8). The structural and accessory proteins contribute to virion structure and assembly, virulence and immune manipulation and invasion (2, 9). However, despite concerted experimental studies, the structural understanding of many of these viral proteins is still missing; for example, in SARS-CoV-2, these include ORF3a, ORF3b, M, ORF6, ORF8, ORF9b, ORF9c, ORF10, and certain parts of ORF1a/b. Here, we utilize a domain-centric computational strategy to systematically study the function and structure of CoV proteins. In our recent work, we have demonstrated that the mysterious SARS-CoV-2 protein, ORF8, belongs to a novel family of the immunoglobulin fold. We also showed that ORF8 is fast-evolving and its function is likely to disrupt the host immune responses (10). In this study, we present results on the function and evolution of novel ion channel proteins in CoVs and other nidoviruses.

Viral ion channels (viroporins) represent a new functional class of proteins which have been identified in different animal viruses, including human immunodeficiency virus (HIV), hepatitis C virus, and influenza A virus (11). These proteins are shown to facilitate several steps of the viral life cycle from genome replication, viroplasm formation, and virion budding, to viral infection (11). CoVs also code for their own ion channels. The SARS-CoV ORF3a was found to function as a potassium-specific channel promoting virus release (12). Thereafter, several other CoV proteins were shown to display similar ion channel activities, including porcine epidemic diarrhea virus (PEDV) ORF3 (13), hCoV-229E ORF4a (14), and SARS-CoV envelope (E) protein (15, 16). Among them, SARS-CoV ORF3a, PEDV ORF3 and hCoV-229E ORF4a, appear to utilize their three transmembrane (3-TM) region to constitute an ion channel in the form of either a dimer or a tetramer, whereas SARS-CoV E protein with a 1TM region forms an ion channel as a pentamer (15, 16). Recently, a similar ion channel activity was also observed in SARS-CoV-2 ORF3a and its structure was solved (17). This provides an opportunity for us to systematically identify other ion channel proteins in coronavirus and related genomes by using sensitive profile-based homology detection and structural modeling methods. As a result, we have identified several homologous protein families, including ORF5 from MERS-CoV, several proteins from beta-CoVs, and ORF3b from alpha-CoVs. Importantly, we show that the well-known Matrix (M) proteins from CoVs and several other nidoviruses are also homologous to the ORF3a proteins and identify their likely ancestral members of the family. We present evolutionary and structural evidence that they utilize distinct conserved residues to constitute the ion conducting pore in the membrane. Finally, we reconstructed the genome composition changes during the evolution of CoVs and other related viruses which suggests an evolutionarily conserved role of ion channels in viral virion assembly and membrane fusion/budding.

## RESULTS AND DISCUSSION

### Unification of M/ORF3 ion channel families in CoVs and other nidoviruses

The ORF3a has two domains including a leading 3-transmembrane (3-TM) region and a beta sandwich domain. We first identified the viral homologs of ORF3a by conducting iterative sequence searches using PSIBLAST against the NCBI NR database (18). A multiple sequence alignment was conducted to identify the evolutionarily conserved residues, and the majority of them are located within the ion channel pore of the 3-TM structure (S1 Fig). Interestingly, when we used HMM profile-based homology detection against Pfam profiles via HHsearch (19), we found two other coronavirus families as the significant hits: one is coronavirus M protein (Pfam ID: PF01635) and the other is ORF3b from alpha-coronavirus (Pfam ID: PF03053). As many viral proteins are not present in Pfam profiles, we conducted a systematic clustering analysis and selected the representatives, excluding the ORF1a/b, for a series of HHsearch analysis. This revealed three other viral protein families that are related to ORF3a and M families, prototyped by ORF5 of MERS-CoVs, ORF4 of 229E-related bat CoV (NCBI accession number: ALK28794.1) and ORF3 of Eidolon bat coronavirus/Kenya/KY24/2006 (NCBI accession number: ADX59467.1). Our analysis extended these relationships to other nidoviruses beyond coronaviruses leading to the unification of many novel M proteins from fish, reptile, and mammal ToroVs.

The sequence identity between these protein families are between 10-20% (Fig 1) and they share the domain composition of having both 3-TM and a β-sandwich domain. In order to exclude the possibility that their similarity is only due to the presence of 3-TM, we further used their β-sandwich domains to conduct the HHsearch analysis, which also revealed comparable results. Fig 1 presents an overall pattern of these viral proteins and their inter-relationship identified by both sequence-profile and profile-profile searches. Each family segregates as a dense sub-network, while the families themselves are linked together by connections revealed mostly via profile-profile analysis. Hence, we term these unified protein families the M/ORF3 superfamily.

**Fig 1.**
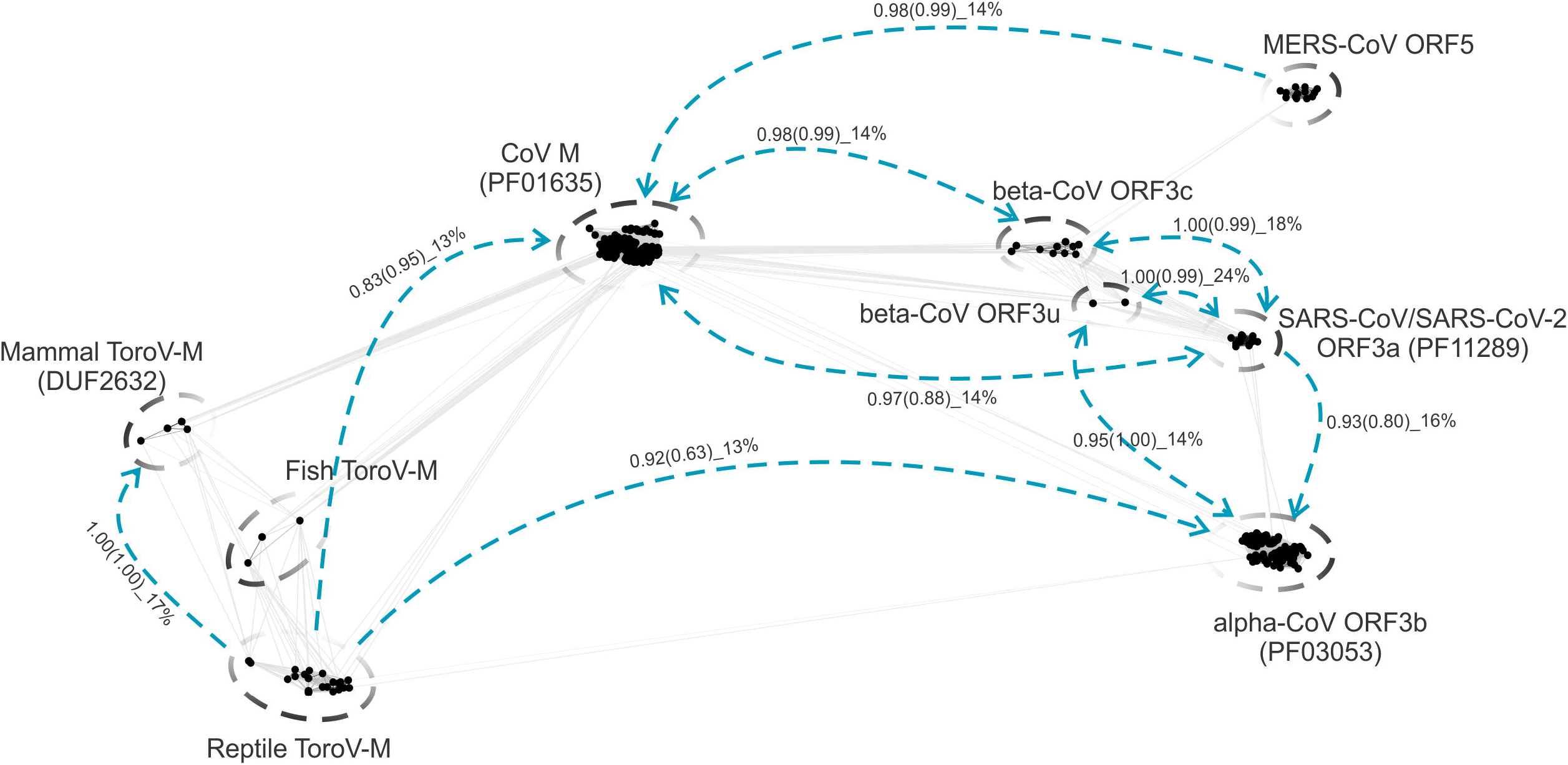
Sequence space of the divergent M/ORF3 proteins of CoVs and ToroVs. Each dot corresponds to a protein sequence. Straight lines indicate the similarity that can be detected by local BLASTP comparisons (e-value cutoff: 0.02). Gapped circles indicate individual protein families. Blue dashed lines indicate the remote relationship detected by the HHsearch program where the arrow indicates the search direction. The probabilities of the HHsearch comparisons with full-length protein sequences and C-terminal β-sandwiches (in brackets) are indicated, followed by the percentage sequence identity.

### Sequence and structural features of the divergent M/ORF3 proteins

Based on both sequence searches and clustering analysis, we divided these proteins into nine families (Fig 1). All of them share a common domain architecture with the 3-TM region followed by a β-sandwich domain. We carefully generated a super-alignment for the β-sandwich domains of these proteins which revealed a comparable eight beta-sheet arrangement despite their low sequence identity (Fig 2). There are no universally conserved polar residues in these C-terminal β-sandwich domains indicating they are not enzymatic modules.

**Fig 2.**
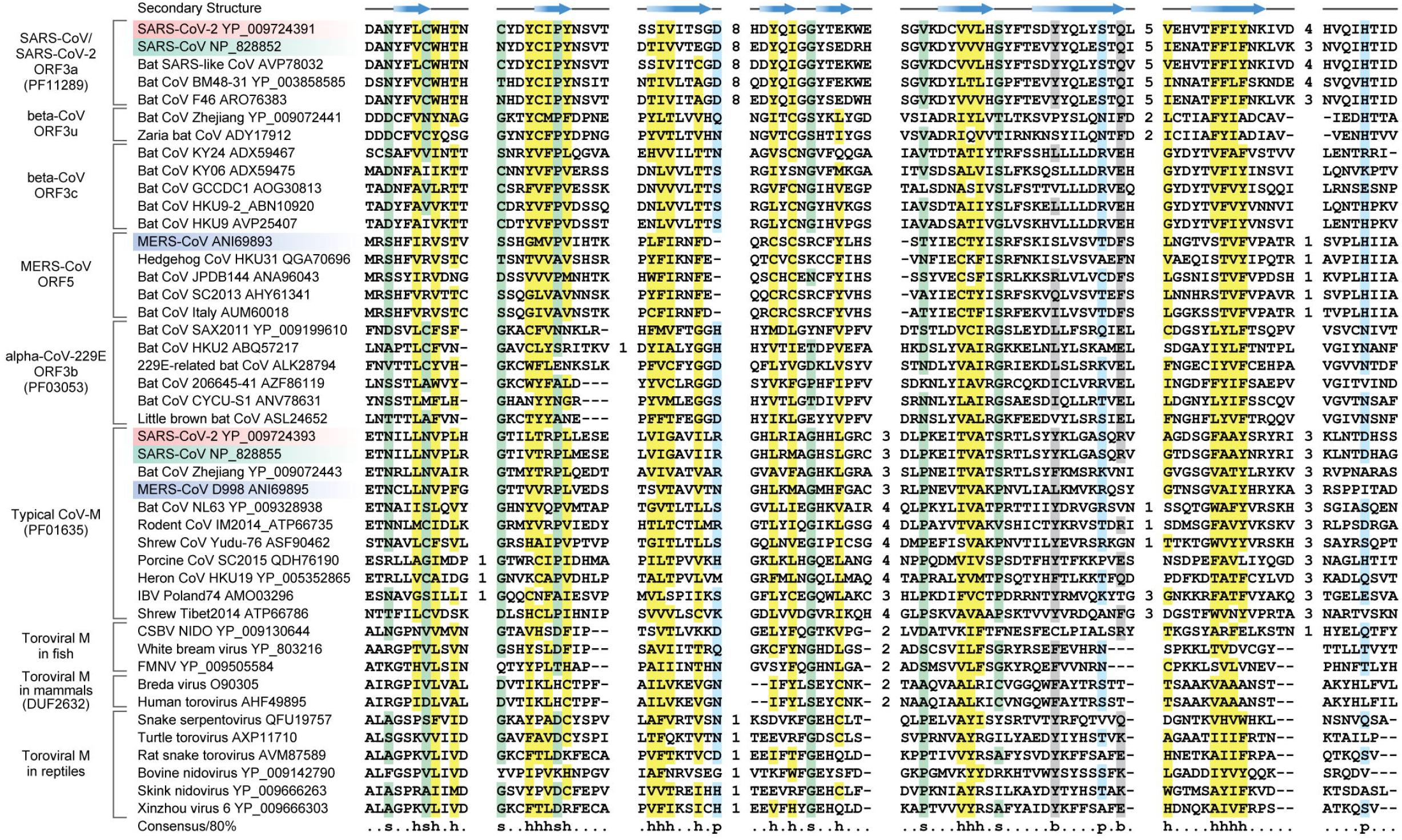
Multiple sequence alignment (MSA) of β-sandwich domains of nine M/ORF3 protein families. The secondary structure is shown above the alignment and the consensus is shown below the alignment, where h stands for hydrophobic residues, s for small residues, b for big residues, and p for polar residues. The numbers are indicative of the excluded residues from sequences. Sequences from three human-infected CoVs are highlighted (SARS-CoV-2 in red, SARS-CoV in green and MERS-CoV in blue)

As ORF3a of the SARS-CoV clade functions as an ion channel (12, 17), we next asked if other viral protein families might have similar functions. We generated homology dimeric models for several representatives of other families (Fig 3) by using the SARS-CoV-2 ORF3a structure as the template (PDB: 6XDC) (17). We then identified the evolutionarily conserved residues for each viral family and evaluated them for any conserved compositional trends. These viral families do not share any conserved residues across the whole superfamily (S2-S7 Figs).

**Fig 3.**
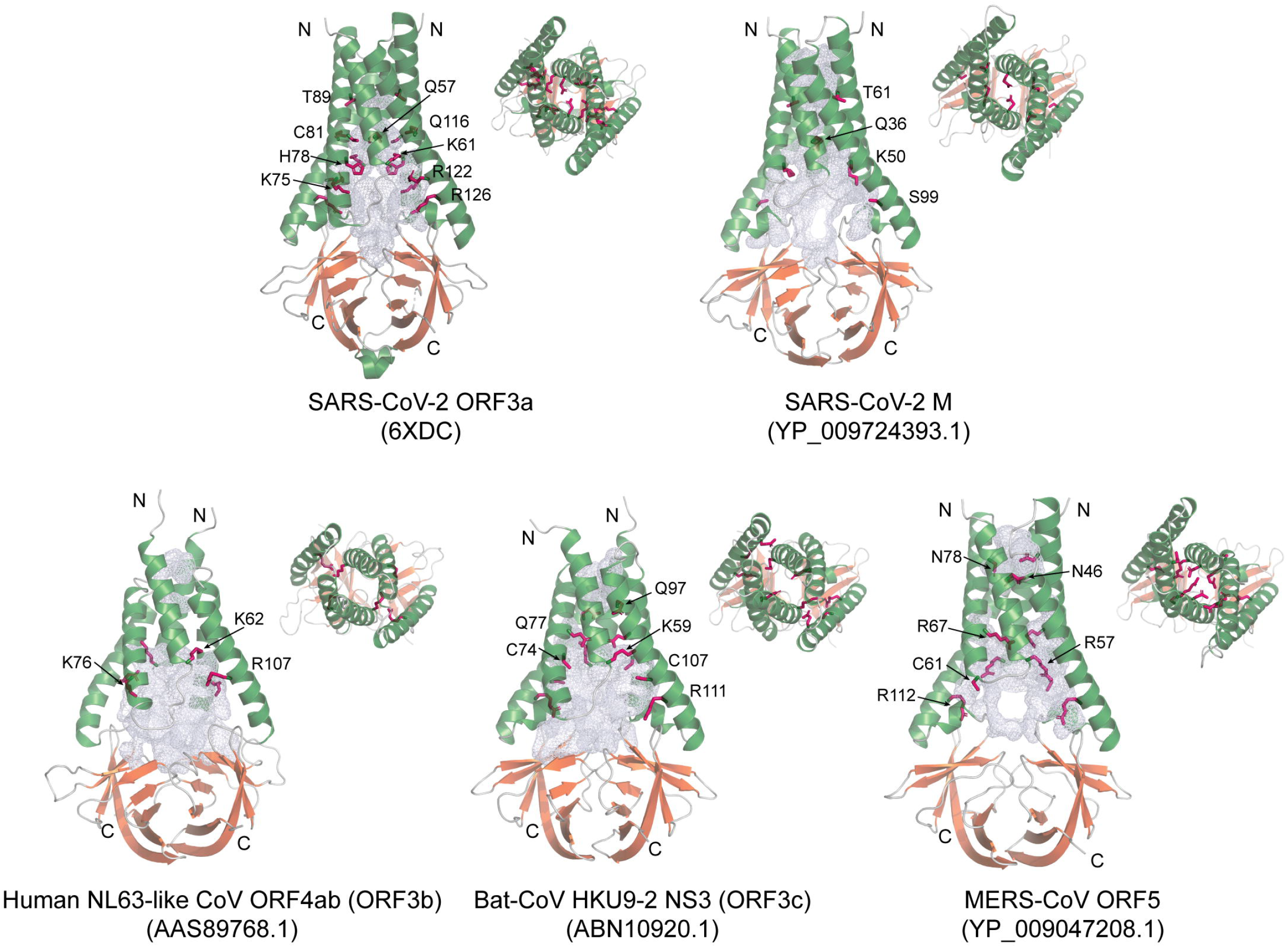
Dimeric structures of SARS-CoV-2 ORF3a and homology models of SARS-CoV-2 M, human-NL63-like-CoV ORF3b, bat-CoV HKU9-2 ORF3c, and MERS-CoV ORF5. Both the side-view and top-view are presented. Alpha-helices are colored in green, β-sheets of the β-sandwich domains in orange, and loops in grey. The channel cavities are colored in grey. The polar residues which are conserved in each family are located on the cavity surface, shown in a purple stick model. The PDB and NCBI ids are shown in brackets.

However, they display a unique conservation of family-specific polar residues, mostly basic residues, which are located in the internal surface of ion channel cavity (Fig 3). This indicates that these residues have been conserved to maintain an aqueous pore within the membrane-spanning region. Therefore, all the novel viral families are potential ion channels whose predicted pore-forming regions are defined by family-specific polar residues.

### Evolutionary relationship of divergent M/ORF3 families

We next sought to examine the evolutionary relationship of these M/ORF3 proteins. Based on the version of the super-alignment of the β-sandwich domains, we conducted phylogenetic analyses using robust Maximum Likelihood and Bayesian inference methods (20-22). The relationship of M/ORF3 families parallel the relationship of viruses that contain them (Fig 4A). The M protein families from ToroVs are clustered together as a firm clade. They form as a sister group of all coronavirus families. Among these CoV families, the CoV-M protein family forms a distinct clade which can be further divided into several subfamilies in Delta-, Gamma-, Alpha-, Beta-CoVs and a new unclassified group typified by a CoV from Guangdong Chinese water skink (NCBI accession number: AVM87576; Fig 4A). All other CoV families form a second major clade, suggesting that they likely share a common ancestor. Among them, ORF3b is only present in alpha-CoVs whereas ORF3c, MERS-CoV ORF5, ORF3u and SARS-CoV/SARS-CoV-2 ORF3a are present in different beta-CoV subgroups. Collectively, due to their relationship with CoV M proteins, we propose to name these divergent ORF3a-like proteins as CoV Matrix 2 (M2) proteins while the typical CoV M proteins as M1.

**Fig 4.**
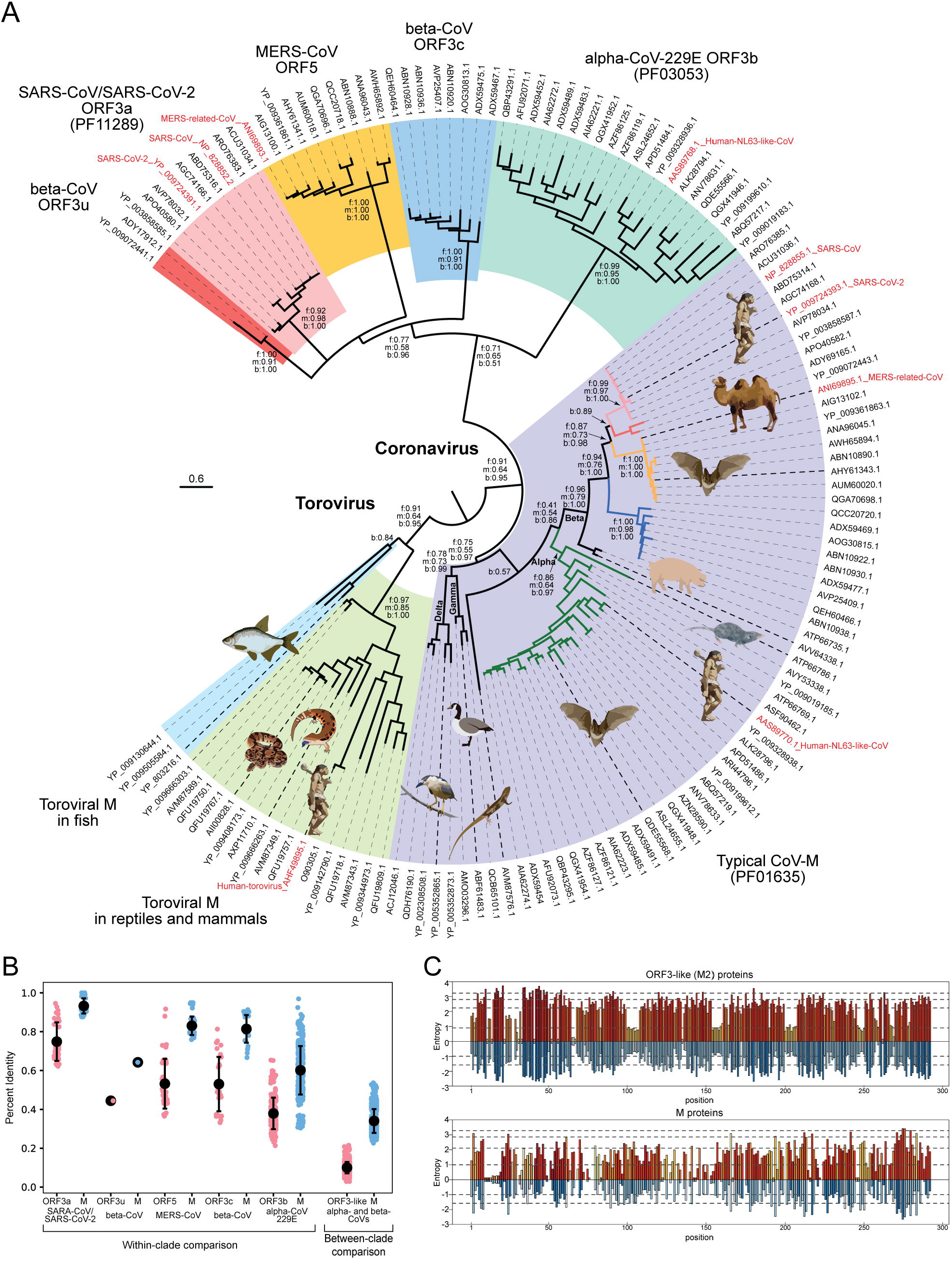
**(A) Evolutionary relationship of M/ORF3 ion channel families of CoVs and ToroVs.** The ML tree with the highest log likelihood (−16934.18) is shown with supporting values for the major branches from three phylogenetic methods: f for FastTree, m for MEGA-ML and b for BI posterior values. Sequences are represented by the NCBI accession IDs and those from human viruses are further labeled with the virus name in red. The cartoon of several viral hosts is shown; its correspondence with the sequence is indicated by the bold dashed lines. Each family is highlighted in a different background. For the sequence within the M family, if they are coupled with one of the ORF3 families, their branches are colored with the same theme, accordingly. **(B) Percent identity comparison between the coupled M and ORF3-like (M2) proteins within different viral clades and between different clades. (C) Shannon entropy plot of the M and ORF3-like (M2) proteins.** Shannon entropy data computed based on regular amino acid alphabet (20 amino acids) are shown above the zero line in shades of orange. Shannon entropy data computed based on a reduced alphabet of 8 residues are shown below the zero line in shades of blue. Where a position shows high entropy in both alphabets, it is a sign of potential positive selection at those positions for amino acids of different chemical characters.

### The coupled ORF3-like (M2) and M proteins display different evolutionary rates

When we examined the genome organizations of the viruses which contain the M and ORF3-like proteins, we found all ORF3-like proteins are strictly coupled with one typical M protein on the viral genome in an order of ORF3-E-M (Fig 5), where ORF3 represents one of the divergent ORF3-like (M2) families, E is the envelope protein, and M is the typical matrix protein of CoVs. The inter-relationships of the alpha- and beta-CoV ORF3-like (M2) families is consistent with the internal relationship of the coupled M proteins on the same viral genome (Fig 4A). This indicates that both the ORF3-like (M2) and M proteins have been vertically inherited from the common ancestor of the alpha- and beta-CoVs.

**Fig 5.**
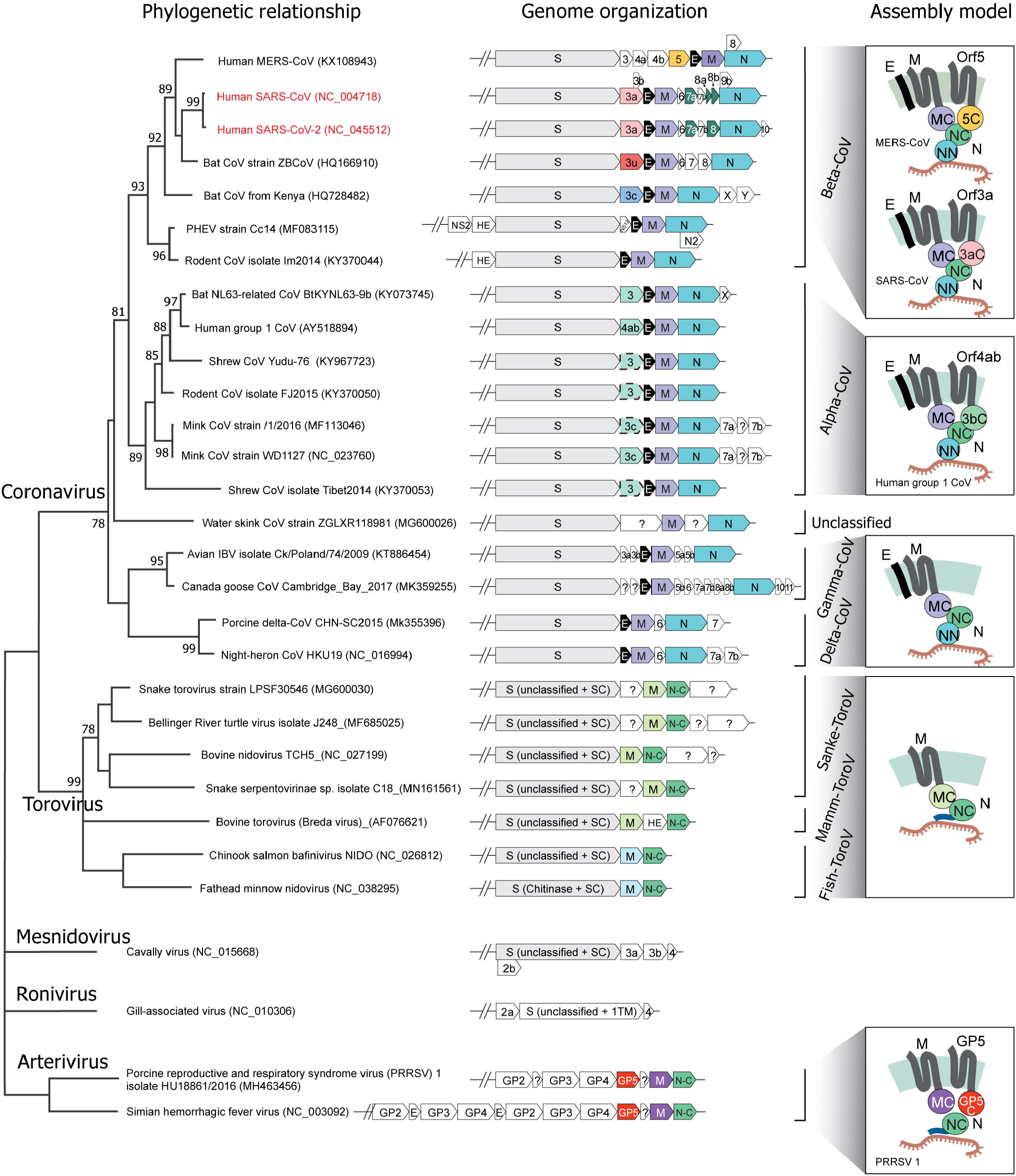
Genomic organization of representative CoVs, ToroVs and other related nidoviruses. The detailed species tree of the CoV-ToroV clade was adapted from the Fig 4. The genomic regions encoding for structural and accessory proteins are illustrated right to the terminal nodes of the phylogenetic tree. The gene names are shown according to their NCBI genome annotation, but the color of the gene box corresponds to different protein families: all CoV Spike proteins in grey, SARS-CoV/SARS-CoV-2 ORF3a in pink, beta-CoV ORF3u in red, MERS-CoV ORF5 in yellow, beta-CoV ORF3c in sky-blue, alpha-CoV-229E ORF3b in pale-green, typical CoV-M proteins in light-purple, reptile and mammal ToroV M protein in light-green, fish ToroV M proteins in light-blue, envelope (E) protein in black, SARS-CoV ORF7a-Ig and ORF8-Ig in dark-green, nucleocapsid (N) protein in blue-green, nucleocapsid (N) protein C-terminal dimerization domain in green, Arterivirus ORF3-like GP5 protein in vermilion and Arterivirus M protein in purple. Several previously unannotated alpha-CoV ORF3b genes are recovered and highlighted in dashed boxes. The predicted assembly models (without Spike) are shown on the right side of genomic organization. Protein abbreviations used in the model are as follows: MC: M protein C-terminal β-sandwich domain; NN: N protein N-terminal RBD domain; NC: N protein C-terminal dimerization domain; SC: Spike protein C-terminal domain; 5C: MERS-ORF5 C-terminal β-sandwich domain; 3aC: SARS-CoV/SARS-CoV-2 ORF3a C-terminal β-sandwich domain; 3bC: alpha-CoV-229E ORF3b C-terminal β-sandwich domain; GP5C: Arterivirus GP5 protein C-terminal β-sandwich domain.

However, these proteins show striking differences in their evolutionary rates. Within each viral group, the sequence identity between the ORF3-like proteins is always significantly lower than the one of the coupled M proteins (Fig 4B). Further, when we compared proteins between different viral clades, we found the average percent identity of M proteins is about 34.0%, whereas the average percent identity of ORF3-like (M2) proteins is only 10.0% (Fig 4B). This indicates that the ORF3 (M2) protein was evolving much faster than the genomically-linked M protein. To better understand the functional difference between them, we conducted the column-wise Shannon entropy analysis (Fig 4C). We found that across the length of the whole alignment, ORF3-like proteins have significantly higher mean column-wise entropy than the M proteins from the same set of viral genomes (2.22 as opposed to 1.52; p=1.6 × 10^−16^ for the H_0_ of congruent means by t-test). By comparing column-wise entropies in both the 20-amino-acid alphabet and a reduced 8-letter alphabet (where amino acids are grouped based on similar side chain chemistries), we found a similar result (1.51 for ORF3-like proteins as opposed to 0.92 for the M proteins; p= 2.2 × 10^−7^). This also indicates that there is a much greater tendency in the ORF3-like (M2) proteins to contain a position that differ drastically in side chain chemistry (e.g. substitution of a charged for a hydrophobic residue) and supports selection for diversification in these proteins as opposed to their M counterparts. This is also supported by the computation of the normalized Kulback-Leibler entropy (S8 and S9 Figs), where positions with strongly negative values indicate a diversifying pressure as opposed to strongly positive values which suggest preservation of conserved configurations. Former positions are significantly more common in the ORF3-like (M2) proteins as opposed to the M-protein (Kruskal-Wallis test, p= 10^−12^).

In conclusion these results suggest that the M proteins are undergoing purifying selection and might have been shielded to a degree from attacks by the host immune system, whereas the ORF3-like proteins are likely to be at the interface of the interactions between the host and virus.

### Evolution of virion assembly mode of CoVs, ToroVs, and other related nidoviruses

Other than M and ORF3-like (M2) proteins, several other proteins are also major structural components of coronavirus, including S, which is a major structural component of the virion involved in cellular receptor binding (23), E, which is a 1-TM ion channel protein critical for envelope formation and membrane budding (24), and N which is essential for viral genome packaging and virion assembly (25, 26). Hence, we investigated if they also show a pattern similar to that of the M/ORF3 proteins. We conducted a genomic composition analysis by using both similarity-based clustering and domain analysis for CoVs, ToroVs, and other known nidoviruses such as roniviruses, mesnidoviruses and arteriviruses (Fig 5).

Consequently, we found that the Spike proteins of three major lineages of nidoviruses such as CoVs, ToroVs, and mesnidoviruses show a conservation at their C-terminal regions (SC; Pfam domain: PF01601) while their N-terminal regions display variations in which the CoV-Spike contains the characteristic NTD and RBD for receptor binding whereas other viral Spikes contain apparently-unrelated regions (Fig 5). The M proteins from both CoVs and ToroVs share a similar architecture of 3TM and β-sandwich domains. Interestingly, we found that Arterivirus contains two proteins with distinct 3TM domains, so-called M and GP5 (S10 and S11 Figs). Both of their N-terminal 3TM domains share similarity and conserved polar residues with the M/ORF3-3TM domain, but their C-terminal domain is an unrelated shorter domain with just 6 β-strands (S10 and S11 Figs). This indicates that the 3TM-type ion channels of the CoVs, ToroVs, and Arterviruses share a common ancestry, but their coupled β-strand-rich C-terminal domains might have different origins. In the case of the N protein, its C-terminal dimerization domain (N-C in Fig 5) shows a wide distribution among ToroVs and Arteriviruses, in addition to CoVs (S12 Fig). These proteins do not possess the RNA-binding domain (RBD) of CoV-N; instead they carry a long N-terminal region which contain a stretch of basic residues (S12 Fig); therefore, we predict that this region functions equivalently in binding negatively-charged RNA viral genome. The E protein and RBD of CoV N are found in all CoVs sharing the common gene order of S-E-M-N. This genomic organization was further modified in alpha- and beta-CoVs which acquired an ORF3-like (M2) gene between the S and E genes. It is likely that ORF3 was derived from a duplication event of M at the ancestor of alpha- and beta-CoVs. Thus, the genome composition analysis helps reconstruct the series of evolutionary events that resulted in the current complex form of the structural proteins of CoVs and other nidovirus (Fig 5).

Other than this buildup, we also observed additional lineage-specific genes. For example, the SARS-related clade has several new genes including ORF6, ORF7 and ORF8 which were inserted between M and N genes (10), while the MERS-CoVs, which have several genes were inserted between Spike and ORF5 genes. They could be newly introduced structural components during evolution.

### FINAL REMARKS

Thus, we have systematically unified and classified many divergent ion channel proteins across several major lineages of nidoviruses, including CoVs, ToroVs and Arterivirus. We show that the core virion component, M proteins of CoVs and ToroVs, share a similar architecture and utilize conserved polar residues to form an ion channel in the membrane. Viral ion channels have earlier been extensively studied in the influenza virus – the transcript of the M1 gene of that virus is alternatively spliced in nuclear speckles to give rise to the mRNA M2 that codes for a proton channel needed for acidification and release of viral ribonucleoproteins in the endosome during invasion (27). This, together with the current observations, suggests that establishment of ionic gradients are important for viral function. It is conceivable that different representatives of the nidoviral superfamily unified in this study provide a diversified set of ion-conductance activities in functions including virion assembly and release.

Our results raise several questions for future research on these viruses: 1) Given that M is the core structural component of virion, does its predicted ion channel activity contribute to virion maturation? 2) What might be the effect of the interactions between the M, ORF3a, E, N and Spike on the predicted conductance of the channels and virion assembly; 3) What are the roles of ORF3 in terms of its interface with host proteins? The final question is also related to the roles of the genetic differences in the host factors resulting in widely different disease outcomes observed for COVID-19.

## MATERIALS AND METHODS

### Protein sequence searches and analysis

To collect protein homologs, iterative sequence profile searches were performed using the programs PSI-BLAST (Position-Specific Iterated BLAST) (18) and JACKHMMER (28), which searched against the non-redundant (nr) protein database of NCBI with a cut-off e-value of 0.005 serving as the significance threshold. Similarity-based clustering was conducted by BLASTCLUST, a BLAST score-based single-linkage clustering method (ftp://ftp.ncbi.nih.gov/blast/documents/blastclust.html). Multiple sequence alignments were built using the KALIGN (29), MUSCLE (30) and PROMALS3D (31) programs, followed by careful manual adjustments based on the profile–profile alignment and predicted structural information. Sequence-profile and profile-profile comparisons were conducted using the HHsearch program (19). Secondary structure was predicted using the JPRED program (32). The consensus of the alignment was calculated using a custom Perl script. The alignments were visualized using CHROMA program (33) and further modified using adobe illustrator. The transmembrane regions were predicted using the TMHMM Server v. 2.0 (34).

### Protein structure prediction and analysis

The Modeller9v1 program (35) was utilized for homology modeling of the structures of SARS-CoV-2 M protein, MERS-CoV ORF5, human NL63-like-CoV ORF3b and Bat-CoV HKU9-2 ORF3c by using the SARS-CoV ORF3a (6xdc) as a template. The dimeric status was modeled according to the template.

The sequence identity between the template and the targets is very low, from 15% between the template and the SARS-CoV-2 M protein, 19% with the MERS-CoV ORF5, 15% with the NL63-like-CoV ORF3b, to 22% with the Bat-CoV HKU9-2 ORF3c. Since in these low sequence-identity cases, sequence alignment is the most important factor affecting the quality of the model (36), alignments used in this analysis have been carefully built and cross-validated based on the information from HHsearch and edited manually using the secondary structure information. For each protein, we generated five models and selected the one that had the highest model accuracy p-value (ranging from 0.06 to 0.013) and global model quality score (ranging from 0.34 to 0.38) as assessed by ModFOLD6 online server (37). Structural analysis and comparison were conducted using the molecular visualization program PyMOL (38).

### Molecular phylogenetic analysis

Based on the super-alignment of the β-sandwich domains of nine M/ORF3 families, we conducted phylogenetic analysis using three robust methods, including the Maximum Likelihood (ML) analysis implemented in the MEGA7 program (21), an approximately-maximum-likelihood method implemented in the FastTree 2.1 program (20), and Bayesian Inference implemented in the BEAST 1.8.3 program (22). For ML analysis, initial tree(s) for the heuristic search were obtained automatically by applying Neighbor-Join and BioNJ algorithms to a matrix of pairwise distances estimated using a JTT model, and then selecting the topology with superior log likelihood value. A discrete Gamma distribution was used to model evolutionary rate differences among sites (4 categories (+G, parameter = 5.2604)). The rate variation model allowed for some sites to be evolutionarily invariable ([+I], 4.41% sites). A bootstrap analysis with 100 repetitions was performed to assess the significance of phylogenetic grouping. For FastTree analysis, default parameters were applied, which include the WAG evolutionary model and the discrete gamma model with 20 rate categories. For Bayesian inference, a JTT amino acid substitution model with a discrete Gamma distribution (four rate categories) was used to model evolutionary rate differences among sites. Markov chain Monte Carlo (MCMC) duplicate runs of 10 million states each, sampling every 10,000 steps was computed. Logs of MCMC runs were examined using Tracer 1.7.1. Burn-ins were set to be 4% of iterations.

The tree with the highest log likelihood from the ML analysis was visualized using the FigTree 1.4.4 program (http://tree.bio.ed.ac.uk/). The bootstrapping values and Posterior values are shown next to the branches.

### Entropy analysis

Position-wise Shannon entropy (H) for a given multiple sequence alignment was calculated using the equation:

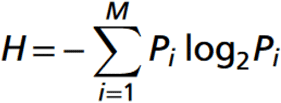

*P* is the fraction of residues of amino acid type *i*, and *M* is the number of amino acid types. The Shannon entropy for the *i*th position in the alignment ranges from 0 (only one residue at that position) to 4.32 (all 20 residues equally represented at that position) in a 20 letter alphabet. Kullback-Leibler divergence (or relative entropy) was computed as described in (39), then centered by the mean and normalized by the range to identify the functionally constrained and diverging positions. Analysis of the entropy values which were thus derived was performed using the R language.

### Genome composition analysis

Open reading frames of viral genomes used in this study were extracted from NCBI GenBank files (40). Protein sequences were subjected to similarity-based clustering by BLASTCLUST with -S at 0.4 and -L at 0.4. Protein clusters were further annotated with conserved domains which are identified by the hmmscan searching against Pfam (19, 41) and our own curated profiles. For previously unknown domains, we used sequence searches followed by multiple sequence alignment and further sequence-profile searches to study their sequence and structural features.

## ACKNOWLEDGEMENTS

Y.T., T. S., and D. Z. are supported by the Saint Louis University start-up fund and the Research Growth Fund – COVID-19 Rapid Response Award. L.A. is supported by the Intramural Research Program of the NIH, National Library of Medicine. P.K.S and M.B.C are supported by NIGMS-5R01GM127783.

## SUPPLEMENTAL MATERIAL

Supplemental material is available online only.

Fig S1. Multiple sequence alignment of SARS-CoV/SARS/CoV-2 ORF3a (PF11289) family

Fig S2. Multiple sequence alignment of Beta-CoV ORF3c family

Fig S3. Multiple sequence alignment of MERS-CoV ORF5 family

Fig S4. Multiple sequence alignment of Alpha-CoV-229E ORF3b (PF03053) family

Fig S5. Multiple sequence alignment of the typical CoV-M (PF01635) family

Fig S6. Multiple sequence alignment of reptile and mammal Torovirus M proteins

Fig S7. Multiple sequence alignment of fish Torovirus M proteins

Fig S8. Relative entropy (Kullback–Leibler divergence) analysis of the M proteins in Alpha- and Beta-CoVs.

Fig S9. Relative entropy (Kullback–Leibler divergence) analysis of the ORF3-like proteins in Alpha- and Beta-CoVs.

Fig S10. Multiple sequence alignment of Arterivirus M proteins

Fig S11. Multiple sequence alignment of Arterivirus Gp5 proteins

Fig S12. Multiple sequence alignment of Nucleocapsid proteins in ToroVs and Arteriviruses.

